# Wild lytic bacteriophages: a potential enhancer of disinfectants’ biofilm inhibition in bio-industrial settings for fruit fly production

**DOI:** 10.1101/2024.03.11.584479

**Authors:** Julio Roberto Matute Torres, Mariandrée Linares Fernández, Pedro Rendón, Pamela Marie Pennington

## Abstract

The pathogenic nature of *Morganella morganii* to fruit flies has been described earlier; its biofilm prevents the larvae of mass reared fruit flies from proceeding to their next life stage by trapping them in their polymeric matrix. For the Mediterranean (*Ceratitis capitata*) fruit fly (“medfly”) mass rearing plants, this can be problem, due to losses generated in the production process. The presence of this bacteria is prevented by using different disinfectants, by themselves, or combined, which can act as selective pressure for the biofilm to resist the disinfectants. In this paper, we propose a method to isolate lytic bacteriophages and use them as quorum quenchers, to prevent microbial consortium from forming biofilms. When tried as viral cocktails, a very significant reduction of biofilm was found. However, when used with one of the two tried industrial disinfectants, the viral effect wasn’t as significant. When interpreted alongside the first experiments, this result opens the door to use bacteriophages to reduce the minimum inhibitory concentration necessary of the disinfectant. A reduction of said concentration would translate into less selective pressure for a resistance against it, and a reduction in costs for the mass rearing process at the time of implementation of the system.

## Introduction

The MOSCAMED “Medfly” program in Guatemala has been very successful since its inception in the late 20^th^ century (1). MOSCAMED was introduced to control Mediterranean fly populations in Guatemala, Belize, and Chiapas, Mexico. This is done through the release of sterile male flies into the environments with wild fly presence. (1;2;3) MOSCAMED breeds *Ceratitis capitata* which, at its larval stage, tends to jump as it reaches the appropriate larval instar (1;2;3). This physiological behavior makes the larva susceptible to bacteria that produces biofilms, since the biofilm matrix captures the larva and prevent it from jumping, and consequently from developing into pupae; -thus becoming a pathogen for the fly. Such is the case of *Morganella morganii*, which produces a biofilm strong enough to trap the larvae in it, killing them. The exact mechanism in which this happens is still unknown, however the lethality of this bacterium to the fly, has been documented by Salas *et al*., 2017, (4). This can severely hinder production efforts.

One of the most effective tools in inhibiting and penetrating biofilms are bacteria’s natural predators: bacteriophages (5;6;7). There are reports that phages have enzymes such as EPS depolymerase, that are capable of degrading the biofilm matrix, however, they tend to be highly specific (6;7) These are usually found within viruses that infect bacteria that produce biofilms, as such, it is logical to think that wild strains of lytic phages that infect *M. morganii* will encode these proteins (4;6;7).

The main objective of this study was to find such phages and test their capabilities of inhibiting the formation of *M. morganii* biofilms, collating the usage of disinfectants regularly used in rearing operations. This would allow to understand the impact that the industrial application of wild phages would have in the process and permit further studies into utilizing depolymerizing enzymes as a biostatic cleaning agent.

## Experimental Procedures

### Bacterial strain

The *M. morganii* strains used were provided by the MOSCAMED rearing facility from larvae-hatching facilities at its campus. A second strain was provided by the Universidad del Valle’s collection and used as a control throughout the experiment.

### Bacteriophage isolation, purification, and enrichment

Wild type bacteriophages were isolated from the medfly egg suspension (greywater) of *C. capitata*’s container collected from MOSCAMED Plant at Santa Rosa, Guatemala. Phage isolation was done as described by Gradaschi *et al*., 2021 (8), with the following modifications: 1mL of logarithmic phase bacteria (OD600 0.4) and 9 mL of sampling water were added to 1mL of 10X Luria-Bertani (LB) broth enriched with CaCl2 10mM and MgSO4 10mM. The solution was incubated for 2h at 37°C, and 120 rpm. Afterwards, 200μL of chloroform were added and stored at 4°C. The resultant lysate was then filtered through a 0.22μm sterile membrane. Phage titre was determined through a spot test (Mirzaei, 2015) using 0.7% LB agarose (45°C), 10 mM CaCl2 and 10 mM MgSO4, logarithmic phase bacteria (OD600 0.4) and the diluted suspension.

Phage purification was done by taking an aliquot with a 10μL tip of a single clear plaque and placed in 3mL of phage buffer (10 mM Tris, pH 7.5, 10 mM MgCl2, 68 mM NaCl) with 50 µL of chloroform. Afterwards, a spot test was performed using 100μl of serial dilutions with phage buffer (8). The procedure was repeated until homogeneous lysis plaques were obtained. Enrichment of the purified phages was performed by adding 1mL of logarithmic bacterial broth and 15mL of LB broth enriched with CaCl2 and MgSO4 to 150μL of the phage suspension. The mix was incubated overnight at 37°C under constant agitation of 120 rpm. The bacterial density was quantified by spectrophotometry. The culture was centrifuged at 6000rpm for 5min and filtered through a 0.22 µm diameter membrane.

### One-step death curve

Death curves were prepared based on Chen *et al*., 2018 (9) using LB broth. Upon reaching an OD_600_ of 0.4, 20mL of bacteria were distributed in a 50mL tube. 200µL of isolated bacteriophage suspension were added to each, working in duplicates. They were incubated at 42°C, and 120rpm. The absorbance was measured, from a 1mL sample, using a Unico 2800 UV/Vis spectrophotometer at 600nm, every 15min.

### Biofilm formation and Minimal Infectivity Concentration (MIC) tests

Optimal conditions for biofilm formation were determined using a 96-well microplate filled with a variable amount of logarithmic bacterial culture in quadruplicate. It was incubated overnight at 37°C without agitation. The microplate was washed with low water pressure and prepared for a crystal violet assay described before by Shukla, *et al*., 2011 (10). The procedure was performed using Biotek ELx 808 microplate reader spectrophotometer and Gen 5 software for quantification analysis.

The optimal amount of phage suspension to evaluate its infectivity was determined by adding optimum established amounts of bacterial culture (OD600 0.4) and LB broth in a 96-well microplate (8). A variable amount of bacteriophages suspension (20-80 µL) was added to the microplate and then was incubated overnight at 37°C. After incubation, water washes were performed for crystal violet assay as described above. When optimized, phages were mixed into cocktails and then added in quadruplicate to each well with bacterial culture (OD600 0.4) and LB broth. The microplate was incubated overnight at 37°C without shaking. After incubation a crystal violet assay was performed.

### Infectivity test and antiseptics

The interaction between phage cocktails and commercial antiseptics was evaluated using a crystal violet assay. Each well was prepared with optimal quantities of bacterial culture, phage cocktail and Povidone-iodine at 5ppm. The microplate was incubated overnight at 37°C without shaking. After incubation a crystal violet assay was performed, as described previously. The same procedure was performed using Chlorhexidine gluconate as antiseptic with the same concentrations.

#### Data Analysis

Mean values from each treatment were analyzed to determine statistical significance. The values of the cocktails presented normality, constant variance, and independence. This was evaluated with descriptive statistics, histogram graphs, residual graphs, and a Shapiro test for each one. ANOVA tests and a series of Tukey tests determined statistically significance compared to the significant control. Kruskal Wallis test was performed on antiseptic treatments to evaluate non-parametric data means and a Dunn’s Multiple Comparison tests to compare them to the control.

## Results

### Isolation, purification, and enrichment of wild phages

Phage plaque formation was identified by a spot test with the aim of selecting the phages to be purified and enriched later. They were identified as MMф with a variable diameter between 3.7 to 4.8 mm. Spot tests determined the viral titer of the enriched isolated phage samples. A total of 6 phages were evaluated. After enrichment, the purified phages from the lysate MMф were selected and identified as MMф1, MMф2, MMф3, MMф4, MMф5, and MMф6. The relevant UFPs determined for each phage isolate are shown in Table 1. Lysis plaque count in soft agarose spot test assay using serial dilutions up to 1.0E-11. The values of Plate Forming Units (PFU) are shown in log for each isolate, and the average values for each phage isolate is also shown. If the number of lysis plates ≥ 10 it is considered a ± 10% error. (*) Individual values are shown due to lack of data.

**Table 1.** UFP per mL of purified phages.

| Phage | Dilution | # Plaques | log UFP /mL | $\bar{x}$ log UFP/mL |
| --- | --- | --- | --- | --- |
| MMφ1 | 1.00E-02 | 15 | 4.176 | 5.760 |
|  | 1.00E-03 | 7 | 4.845 |  |
|  | 1.00E-04 | 8 | 5.903 |  |
|  | 1.00E-06 | 13 | 8.114 |  |
| MMφ2 | 1.00E-02 | 3 | 3.477 | 4.628 |
|  | 1.00E-04 | 6 | 5.778 |  |
| MMφ3 | 1.00E-03 | 6 | 4.778 | 9.017 |
|  | 1.00E-11 | 18 | 13.255 |  |
| MMφ4 | 1.00E-02 | 5 | 3.699 | 4.625 |
|  | 1.00E-03 | 5 | 4.699 |  |
|  | 1.00E-04 | 3 | 5.477 |  |
| MMφ5 | 1.00E-04 | 2 | 5.301 | 5.301* |

### One-step death curve

A death curve can be seen in Figure 1 for purified wild phages. Linear growth is shown in the first 60 minutes of the control (dashed lines; grey) while bacterial culture with phage treatments: MMф1, MMф2, MMф3, MMф4 and MMф5 show disruption in the linearity of the curve, as well as less absorbance at the end of the evaluation period (continuous line; black). In contrast, treatment with phage MMф6 showed a poorly stable OD_600_ measurement and close to control growth levels; therefore, it was discarded for subsequent infection tests.

**Figure 1.**
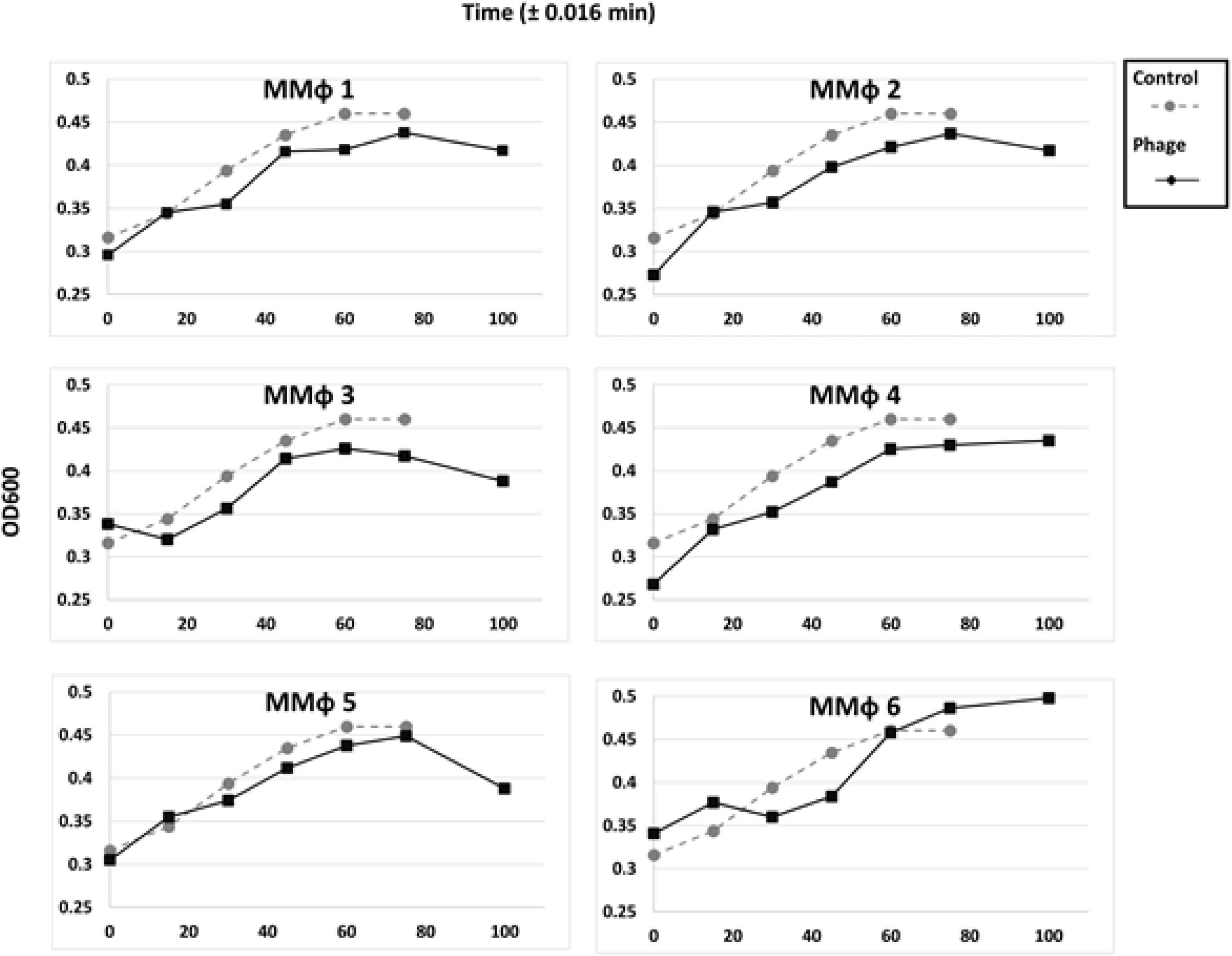
Absorbance according to time at death curve for *M. morganii* with phage treatment. The control (grey): culture of bacteria at the beginning of the logarithmic phase without phage treatment (black).

### Minimal Infectivity Concentration tests

The viral MIC for *M. morganii* was measured adding variable quantities of phages to 160µL of bacteria with an OD_600_ of 4.0. The means of the treatments are shown in Figure 2. Volume treatment of 20-40 µL for MMф1, MMф2 and MMф3 showed no apparent difference against the mean control. However, volumes range from 60 to 80 µL presented greater inhibition; except for phage MMф4 and MMф5 that worked better only with 60 µL. MMф6 was excluded due to its low lethality proven in the death curve procedure.

**Figure 2.**
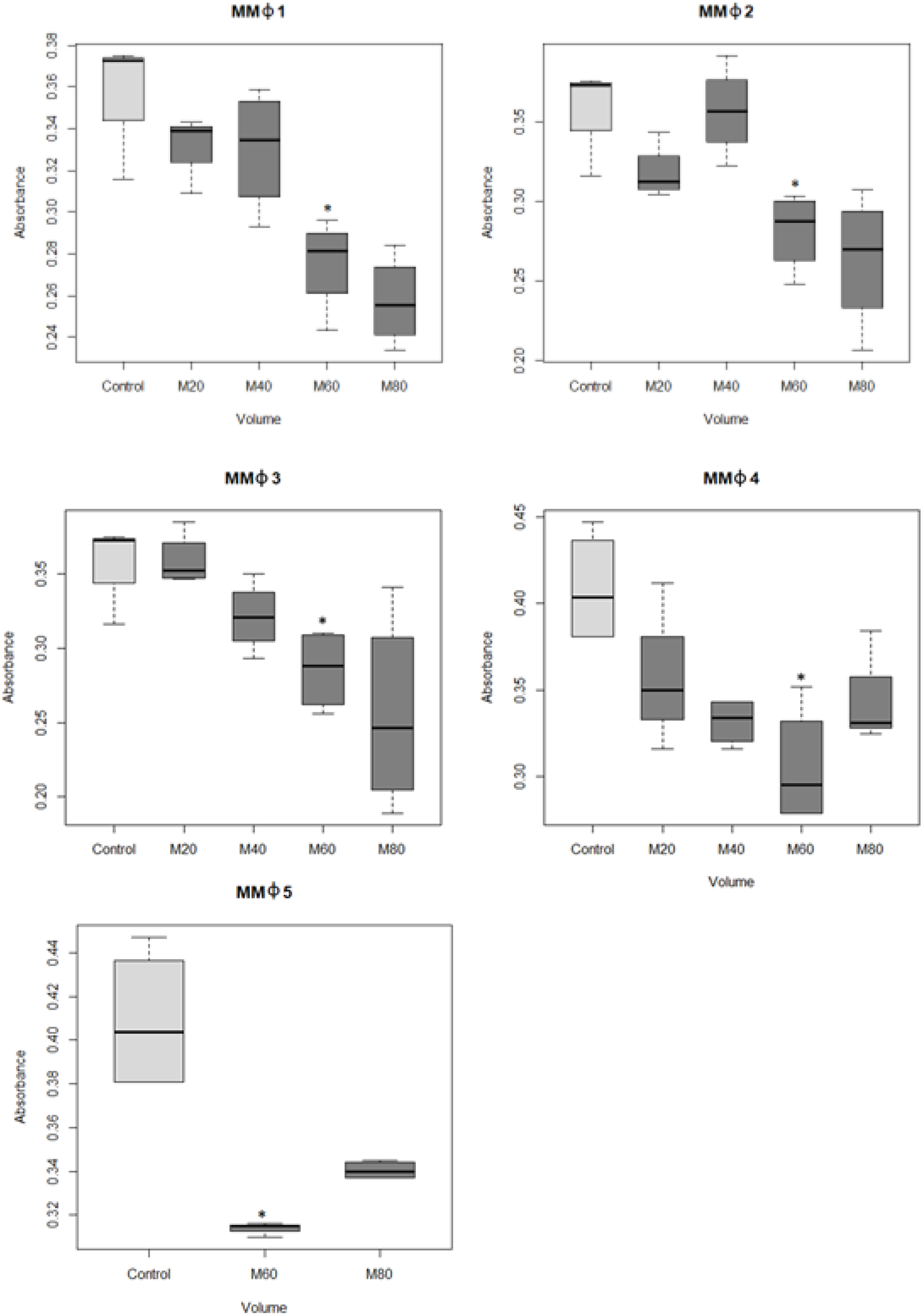
For each phage volume varying from 20-80µL MMф1, MMф2, MMф3, MMф4 and MMф5 were used. In all five cases, adding 60µL of phages (with titers shown in Table 1) proved a significant decrease in bacterial growth.

Every combination of the five wild lytic phages was used to produce a different viral cocktail mixture. The mixtures were co-cultured with *M. morganii* to determine their prowess as biofilm inhibitors, using a crystal violet assay to measure biofilm growth after exposure. The results in Figure 3 (using the mixtures shown in Table 2) show that 5 cocktails hindered bacterial growth in a statistically significant manner (p-value < 0.05). These five cocktails were chosen for the subsequent analysis using disinfectants.

**Table 2.**
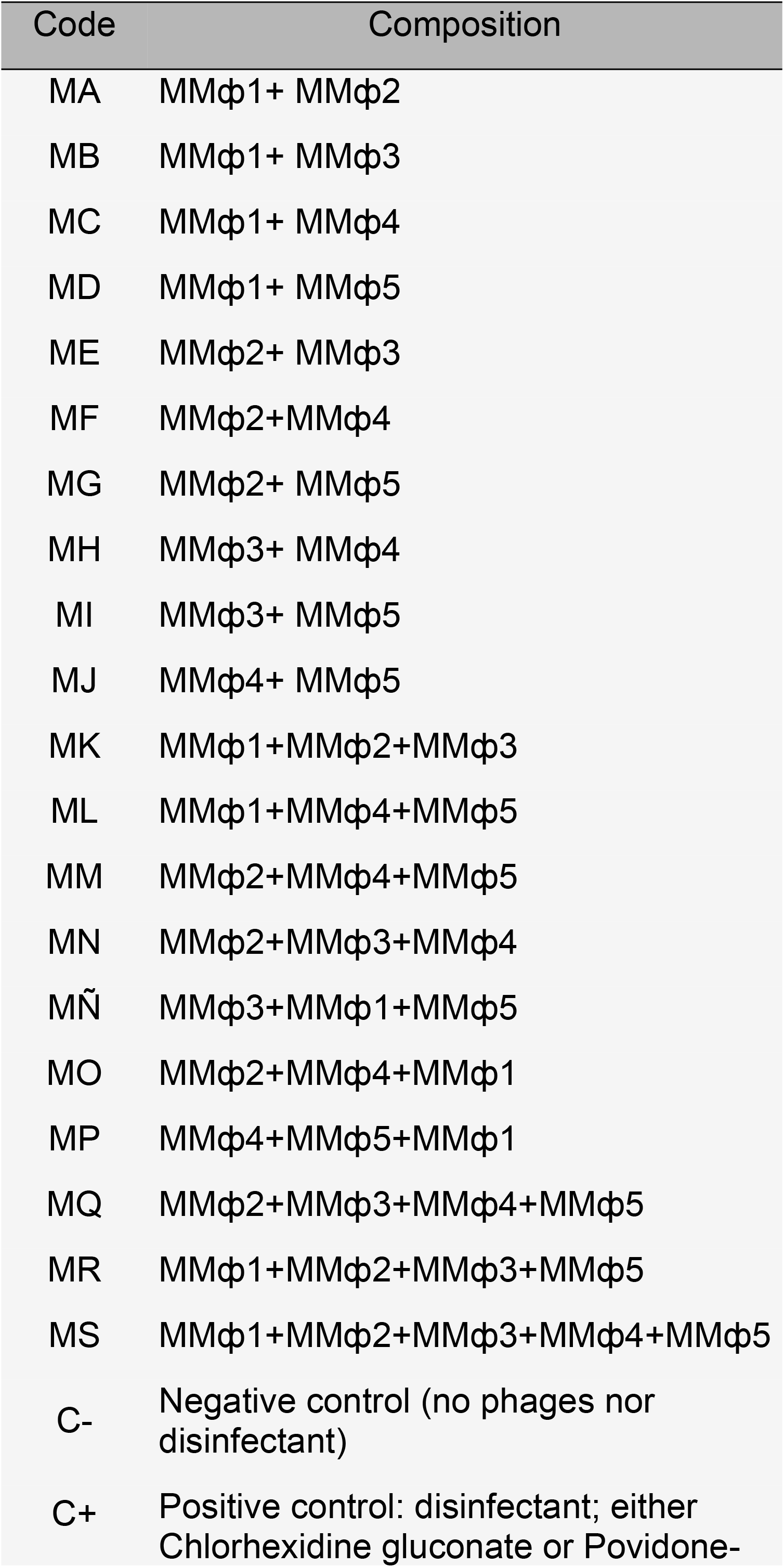
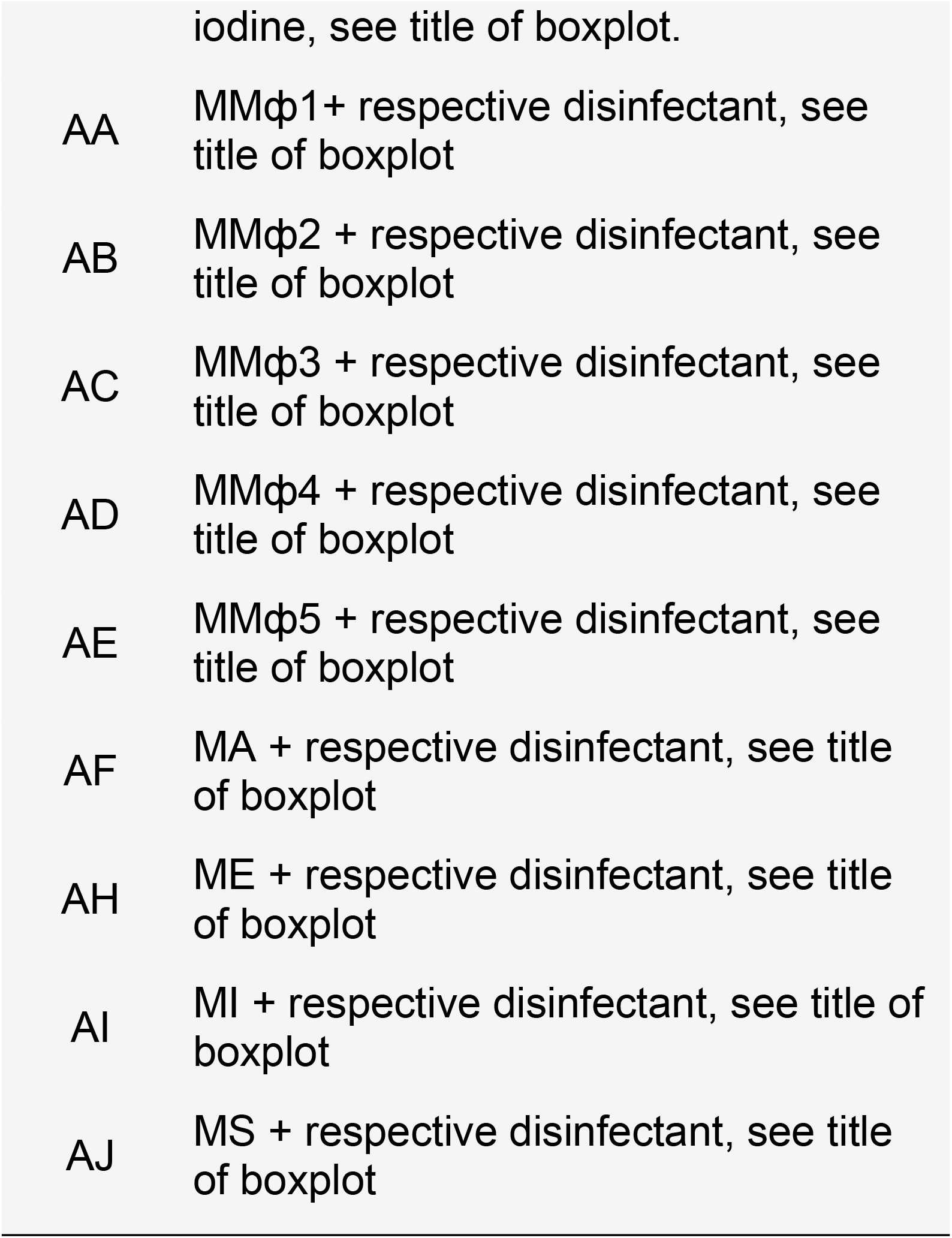
Composition of phage cocktails.

**Figure 3.**
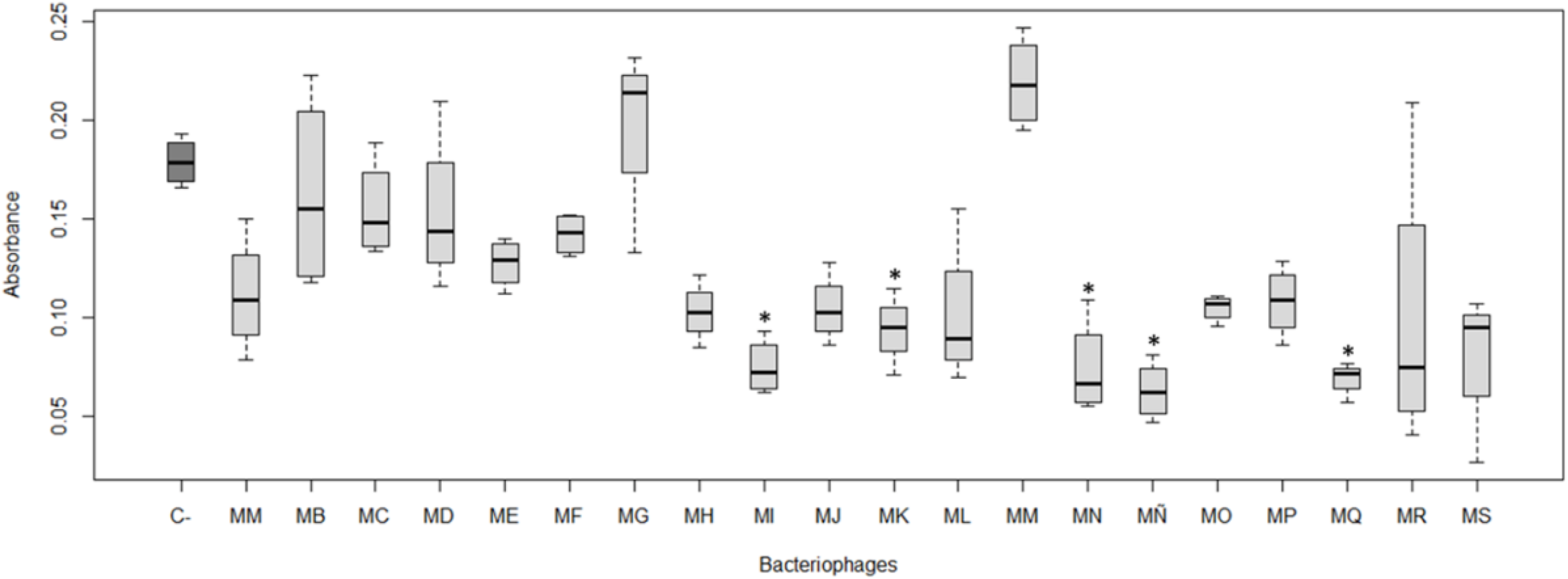
Boxplot of total biofilm formation with phage mixture treatment. C-(dark grey) is the negative control, showing a final absorbance of 0.179 ± 0.010. The following cocktails showed a significant decrease (p ≤ 0.05) compared to the control: MI with 0.073 ± 0.014 (p value 1.3702E-3, MK with 0.094 ± 0.018 (p value 2.46528E-2), MN with 0.074 ± 0.025 (p value 1.2118E-3), MÑ with 0.063 ± 0.015 (p value 1.812 E-4) and MQ with 0.069 ± 0.009 (p value 3.5846 E-3). Composition of the mixtures is indicated in Table 2.

The interaction of phage cocktails with antiseptics for biofilm inhibition was evaluated through a crystal violet assay. Figure 4 shows that every time the disinfectant was supplied with any bacteriophages mix, a decrease in biofilm formation was observed. However, a statistically significant reduction (*p* value < 0.05), was only achieved with mixtures AA, AC, AE, AF, AG, AH, AI and AJ for Chlorhexidine gluconate; and AC, AD, AE, AF, AG, AH and AI for Povidone-iodine against their negative controls. The absorbance of the positive controls (C+) was as expected in Chlorhexidine gluconate (A), though not for Povidoneiodine (B), being higher than that of the negative control. The cocktail treatments showed a larger range of variabilities when used with Povidone-iodine, as seen in Figure 4 (B).

**Figure 4.**
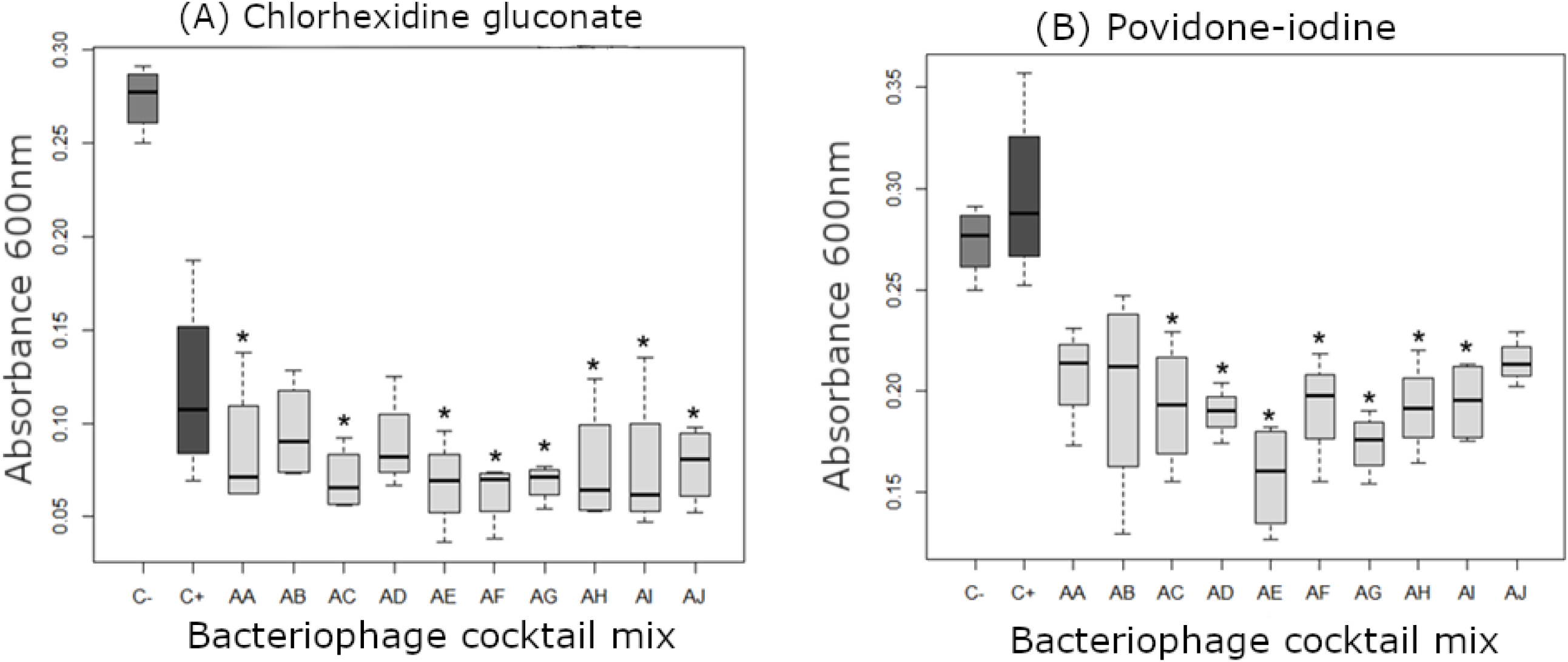
Absorbance of total biofilms with antiseptics and *M. morganii* phage cocktails. (C-) is bacterial culture without change, (C+) bacteria co-cultured with an antiseptic. AA through AJ corresponds to bacterial culture with an antiseptic and a phages cocktail described in Table 2. Significant biofilm inhibition (p ≤ 0.05) is shown in (A) Chlorhexidine gluconate: AA (*p* value 0.022), AC (*p* value 0.005), AE (*p* value 0.003), AF (*p* value 0.002), AG (*p* value 0.004), AH (*p* value 0.005), AI (*p* value 0.002), and AJ (*p* value 0.015); and for (B) Povidoneiodine: AC (*p* value 0.022), AD (*p* value 0.010), AE (*p* value 2.88E-4), AF (*p* value 0.025), AG (*p* value 9.33E-4), AH (*p* value 0.018), and AI (*p* value 0.020).

## Discussion

Bacterial survival has been guaranteed through the development of different mechanisms. Among these, the formation of biofilms has conferred on them pathogenic mechanisms, that both resist antimicrobial substances, and act as a virulence factor. (6;11) In a more specific case, *Morganella morganii*’s biofilm hinders Medfly mass rearing for action programs involving *Ceratitis capitata*, due to the interaction between the fly’s larval stage and the biofilm’s matrix (12). As such, the prevention of this biofilm is of utmost importance to the ongoing success of the initiative. The objective of this study was the analysis and evaluation of the usage of wild-type lytic bacteriophages in conjunction with industrial disinfectants: Povidone-iodine and Chlorhexidine gluconate as biofilm inhibitors. Six specific phages for *M. morganii* were isolated from greywater, five of those were considered apt to be used in treatment, based on their lytic abilities, and apparent infective capacity, which is understood as the capacity of a bacteriophage to enter a host cell and exploit its resources to replicate and produce new viral particles.

To obtain a better read from the plaques formed in point tests, bacteria were grown at 40°C. The added stress allowed for a larger plaque diameter, allowing for an easier analysis. The death curve analysis (Fig. 1) allowed for the lytic capacities of the isolated phages to be compared. This, in turn, was used to select bacteriophages with complementary death curves in the different cocktails that were added to the disinfectants. The reasoning for this lies in the hypothesis that the curvature of the death curves is affected by the metabolic stages of the host. Different viruses will vary their infective capacity at different metabolic stages of the cell; thus, using bacteriophages with different mechanisms will ensure maximum inhibition of biofilm production.

The cocktail composed from MMϕ1, MMϕ3, and MMϕ5 (MÑ, Table 2) presented the largest inhibitory capacity, with the lowest variance, out of all the cocktails tested. On the other hand, the cocktail composed from MMϕ2, MMϕ4, and MMϕ5, (MN, Table 2) allowed for a greater biofilm formation than that of the positive control. This could have been caused by oversaturation of pathogenic pathways. If two or more of those phages shared an entrance mechanism, the competition amongst themselves might not have permitted a proper cellular adsorption, reducing the overall capacity of the cocktail’s infectivity. This, in turn, would have increased the odds of selecting resistant strains multiplied quickly, allowing for a more rapid response from their bacterial anti-phage systems.

It must be noted that *Ceratitis capitata* secretes antimicrobial peptides with a positive cationic charge. These peptides pose a selective pressure for bacterial biofilm to resist positively charged peptides. However, there isn’t a lot of research available on them (12;13), or how these peptides interact with other antiseptic substances.

Povidone-iodine is povodine-iodine. The mechanism used to disinfect is the slow release of negatively charged iodine, which iodizes the cellular lipids, favoring the oxidation of cytoplasmic compounds, triggering cellular death (14;15). The *ceratotoxin* peptides can, then, favor the production of negatively charged biofilms, thus, selecting resistance of bacteria to antimicrobial peptides. However, further research in the interaction between these toxins and biofilm is needed to prove this hypothesis.

The nature of povodine-iodine releases iodine over time, to reduce the damage to eukaryotic cells. However, that timed release could prove enough to allow *M. morganii* to produce a negatively charged biofilm that dramatically reduces the impact of iodine exposure on the bacterial population. Thus, the only inhibitions of biofilm observed were triggered by the bacteriophage cocktails (as shown in Figure 4). A study into the biofilm-specific resistance against this disinfectant would be a valuable addition to prevent usage that selects resistances of disinfectants, as was shown here.

In foresight, a limitation of this study was the absence of a different iodine-based disinfectant as a control to Povidone-iodine. As such, a hypothesis that arises from this experiment is that non-timed release of iodine (such as the exposure to Lugol, for example), will in fact inhibit the formation of *M. morganii* biofilm. However, the increased toxicity to *C. capitata* must be considered before suggesting this to be a workable solution.

The effect of the bacteriophage cocktails when in conjunction with Chlorhexidine gluconate is not significant to that of the Chlorhexidine gluconate acting on its own. This raises the question of whether the viral cocktails can be used to reduce the MIC of an effective disinfectant, in turn, reducing the costs of preventing further *M. morganii* outbreaks.

The mixed results shown for biofilm inhibition in Figure 4 are evidence that bacteriophages can act as quorum quenchers and increase the efficacy of disinfectants as biofilm deterrents. The addition of wild bacteriophages is not cost-intensive when applied correctly. It should also be noted that, if applied, this system should be complemented with quality control that ensures that the bacteriophages are still lytic enough to prevent the biofilm from forming. The characterization of the viruses used would also prevent the spread of lysogenic phages that can easily be a means of horizontal gene transfer for pathogenic isles.

The usage of disinfectants in conjunction with wild lytic bacteriophages can be used as a technique to prevent biofilm formation of *Morganella morganii*. However, further studies are necessary as to which disinfectants are best suited to this task. A thorough characterization of the viruses being used would prove useful, allowing for a more thought-out preparation of the phage cocktails, obtaining better results. A secondary study that uses a lesser concentration of Chlorhexidine gluconate should also be conducted, to evaluate if the MIC is affected with the addition of lytic bacteriophages. The Povidone-iodine results should be confirmed using a different disinfectant based in iodine, without povidone; furthermore, the MIC of povidone would prove incredibly useful to evaluate the disinfectant’s capacity to act as a biofilm inhibitor, with or without the addition of viruses.

## Acknowledgments

This study was conducted as technical collaboration between MOSCAMED Program, the *Centro de Estudios en Biotecnología* and the *Departamento de Bioquímica* of *Universidad del Valle de Guatemala*.

